# The depletion of MARVELD1 leads to placenta accreta via integrin β4-dependent trophoblast cell invasion

**DOI:** 10.1101/106807

**Authors:** Yue Chen, Hui Zhang, Fang Han, Lei Yue, Chunxiao Qiao, Yao Zhang, Peng Dou, Weizhe Liu, Yu Li

**Author notes:** These authors contributed equally to this work (Yue Chen, Hui Zhang).

## Abstract

The mammalian placenta is a remarkable organ. It serves as the interface between the mother and the fetus. Proper invasion of trophoblast cells into the maternal decidua is required for a successful pregnancy. Previous studies have found that the adhesion molecule integrin β4 plays important roles during trophoblast cell invasion. Here, we found that the overall birth rate of the MARVELD1 knockout mouse is much lower than that of the wild-type mouse (*P*<0.001). In E18.5 MARVELD1 knockout mice, we observed an over-invasion of trophoblast cells, and indeed, the pregnant mice had a partial placenta accreta phenotype. The HTR8/SVneo cell line was used as an in vitro model to elucidate the underlying mechanisms of MARVELD1-mediated trophoblast invasion. We detected a diminished expression of integrin β4 upon the downregulation of MARVELD1 and enhanced migration and invasive abilities of trophoblast cells both in vivo and in vitro. The integrin β4 rescue assay also supported the results. In conclusion, this study found that MARVELD1 mediated the invasion of trophoblast cells via regulating the expression of integrin β4.

## Introduction

The mammalian placenta is a remarkable organ. The robust growth of the fetus is dependent on it. It forms the interface between the mother and fetus and is needed for gas and nutrient exchange as well as waste disposal. In addition, the placenta acts as a barrier against the maternal immune system by producing hormones and growth factors(Rossant and Cross, 2001; Watson and Cross, 2005). In humans and mice, the mature placenta is composed of three major layers: the outer layer is called the maternal decidua, which contains decidual cells of the uterus as well as the maternal vasculature that mediates blood exchange with the implantation site. The middle layer acts as a “junctional region”, which is important for the attachment of the placenta to the uterus; it contains trophoblast cells that invade the maternal decidua. The inner layer is called the labyrinth, which is composed of an abundance of branched villi for efficient gas and nutrient exchange(Rossant and Cross, 2001). Although the detailed architecture of the placenta in humans and mice are somewhat different, their overall structures and the molecular mechanisms are quite similar(Rossant and Cross, 2001). Thus, the mouse is an ideal model for studying placental development.

During placenta development in humans, the cytotrophoblast cells from anchor villi differentiate into extravillous trophoblasts (EVTs), lose its proliferative phenotype, and gain an invasive phenotype(Irving et al., 1995), forming three major subtypes: villous cytotrophoblasts (CTBs), syncytiotrophoblasts and EVTs(Fitzgerald et al., 2008; Gude et al., 2004; Lash, 2015). Similar to cancer progression, which involve cellular movement into foreign tissue, trophoblast invasion into the maternal decidua is tightly controlled by many adhesion molecules, including integrins(Damsky et al., 1992; Damsky et al., 1994), matrix metalloproteinases (MMPs) and metalloproteinases inhibitors (TIMPs) (Anacker et al., 2011; Bischof et al., 2000; Godbole et al., 2011). In particular, cytotrophoblast cell invasiveness is associated with different cell surface molecules, which include integrins. Integrins are a family of heterodimeric transmembrane receptors that are formed by two different chains, the α (alpha) and β (beta) subunits, and are expressed on the basal cell surface; it mediates cell adhesion to extracellular matrix (ECM) though the amino acid sequence Arginine-Glycine-Aspartic acid (RGD) (Ruoslahti, 1991; Ruoslahti, 1996). Integrins play important roles in cell adhesion and migration. During the invasion process, the expression of integrin α1β1 and α5β1 are upregulated whereas integrin α6β4 is downregulated (Damsky et al., 1994; Maltepe and Fisher, 2015).

Cellular invasion during placenta development, especially cytotrophoblast differentiation, is a complex process that is tightly regulated. Proper invasion of trophoblast cells into the maternal decidua is required for successful pregnancy(Huppertz, 2008). Disruption of this tightly controlled process can lead to placental deficiencies, resulting in pregnancy complications(Lash, 2015) such as early miscarriage(Hustin et al., 1990), late miscarriage(Ball et al., 2006), preeclampsia(Pijnenborg et al., 1991), fetal growth restriction(Khong et al., 1986), preterm birth(Kim et al., 2003), and placenta accreta(Khong and Robertson, 1987). Placenta accreta is a disorder of human placentation characterized by the abnormal attachment or invasion of placental tissue into the underlying uterine musculature (Dashraath and Lin, 2016; Jauniaux et al., 2016; Luke et al., 1966). Despite the importance of cytotrophoblast cell invasion in placenta development, very little is known about the factors that control this process in vivo.

Nuclear factor MARVELD1 (MARVEL domain-containing 1) is a novel tumor suppressor candidate, which is widely expressed in human normal tissues but is down regulated in multiple carcinomas by promoter methylation (Wang et al., 2009). Previous studies have found that mouse MARVELD1 is a nuclear protein that inhibits cell migration (Zeng et al., 2011). Notably, in non-small cell lung cancer (NSCLC), MARVELD1 regulates the balance of integrin β1/β4, integrin β1/β4-mediated cell surface ultrastructure, and EMT (Yao et al., 2015). Here, we addressed the role of MARVELD1 in mouse placenta development. We demonstrate that the knockout of MARVELD1 in mouse placenta induced the downregulation of integrin β4 and further suppressed integrin β4-mediated cell adhesion; therefore, this novel nuclear factor plays an essential role in the regulation of integrin β4 and associated functions. These molecular mechanism triggers the over-invasion of trophoblast cells, which leads to placenta accreta and a difficult birthing process in *MARVELD1*^-/-^ mice. This study accelerated our efforts in understanding the critical role of the placenta in successful pregnancies.

## Materials and methods

### Cell lines and reagents

Human cell lines NIH/3T3, NCI-H460, MDA-MB-231, NCI-H28, NCI-H2452, HTR-8/SVneo, IRR-MRC-5, and WI38 were purchased from the American Type Culture Collection (ATCC, Manassas, VA). Stable cell lines NIH/3T3-MARVELD1-V5 and NIH/3T3-PC cells were maintained by our lab. Antibodies against MARVELD1 (ab91640), Cytokeratin18 (ab668), and integrin β4 (ab182120) were purchased from Abcam (Cambridge, MA). Antibodies against V5 (#13202) were purchased from Cell Signaling Technology. Antibody against GAPDH (TA-08) was purchased from ZSGB-BIO (ZSGB-BIO, Beijing, China). Laminin (L4544) was purchased from Sigma, TGF-β1 (240-B) was purchased from R&D, Lipofectamine^®^ 2000 transfection reagent (Cat#11668019) and cell tracker (Cat#C34552) were purchased from Invitrogen, TRIzol reagent (Cat#11667165001) and FastStart Universal SYBR Green Master (ROX) (Cat#04913850001) were purchased from Roche, the PrimeScript^TM^RT reagent Kit with gDNA Eraser (Cat#RR047Q) was purchased from Takara, and the Dual-Luciferase reporter assay system (Catalog no. E1910) was purchased from Promega.

### MARVELD1 knockout mice

All animal experiments were performed in strict accordance with the recommendations in the Guide for the Care and Use of Laboratory Animals from the Harbin Institute of Technology. The MARVELD1 knockout mice were generated by Biocytogen (Beijing, CHINA) by injecting conditional MARVELD1 knockout murine ES cells (conditional Cre-loxP target vector cassette) into blastocysts with a C57BL/6 background (C57BL/6J-Tyrc-2J) to obtain chimeric mice. MARVELD1 heterozygous animals were backcrossed to wild-type C57BL/6 mice to obtain conditional MARVELD1 knockout mice without the albino phenotype (Tyrc-2J). Then, the conditional MARVELD1 knockout mice were crossed with Ella-Cre mice to obtain MARVELD1 knockout mice. Mice were genotyped by PCR using primers that detected wild-type and knockout sequences. Primers sequences are available in the Table S1.

### Cell culture and RNA interference

HTR-8/SVneo cells were derived by transfecting the cells that grew from chorionic villi explants of human first-trimester placenta with the gene encoding simian virus 40 large T antigen. These transfected trophoblasts were used for the study of trophoblast biology and placental function. Cells were cultured following the protocol from ATCC (Gibco, NY). Cells were cultured in RPMI-1640 medium with 5% fetal bovine serum at 37 °C in an incubator with 5% CO_2_.

For RNA interference, MARVELD1 siRNAs and scrambled siRNAs were designed by the GenePharma Company. Cells were transfected with Lipofectamine^®^ 2000 Transfection Reagent following the manufacturers’ protocol. The knockdown efficiency of the siRNAs was determined by real-time PCR.

### Reverse transcription and real-time PCR

Total RNA was extracted and purified using the TRIzol reagent. The concentration of each RNA sample was determined by using the NanoDrop 2000 spectrophotometer (Thermo Fisher Scientific Inc.). One µg of total RNA was used as a template for reverse transcription into cDNA using the PrimeScript^TM^RT reagent Kit with gDNA Eraser according to the manufacturer’s instructions. Primers are available in the Table S1.

Quantitative real-time PCR was performed with FastStart Universal SYBR Green Master (ROX) according to the manufacturer’s instructions using the ViiTM 7 Real-time PCR system (Applied Biosystems, Foster City, CA, USA).

### Western blotting

Western blotting was performed as described previously. Briefly, total protein was separated on a precast 12% polyacrylamide gel and blotted with antibodies for integrin β4 (diluted 1:1000), integrin β1 (diluted 1:1000) and GAPDH (diluted 1:20000). Data were analyzed and quantified by using the Odyssey infrared imaging system application software v3.0.

### Histological analysis

For placental tissue, 5 μm sections were used. Sections were collected and fixed in 4% paraformaldehyde, dehydrated, embedded in paraffin and sectioned. Histological sections were stained with hematoxylin and eosin (H&E) or used for other analyses such as immunohistochemistry (IHC) and immunofluorescence (IF) staining assays.

### IHC and IF

Sections were washed three times in PBST (PBS with 0.1% Triton X-100), blocked with 10% normal serum/PBST from the same species as the secondary antibody, and incubated overnight at 4 °C with primary antibodies: Mouse anti-Cytokeratin 18(1:500), Rabbit anti-integrin β4 (1:250), and MARVELD1 (1:100). The primary antibodies were removed, and the sections were washed three times with PBST for 5 minutes each.

For IHC, 100-400 µl of biotinylated secondary antibodies, diluted in TBST per the manufacturer’s recommendation, was added to each section. Then, the antibody staining was developed using established procedures.

For IF, the antibodies were applied in the same manner as for cell staining. The secondary antibodies were from Invitrogen. Stained sections were visualized using a Zeiss AXIO Zoom. V16 Stereo Zoom Microscope.

For cell staining, cells were fixed in 4% paraformaldehyde for 15 minutes on ice, blocked with 5% normal serum from the same species as the secondary antibody, and stained using established procedures. Trophoblast cells were detected using anti Cytokeratin18 (1:500) and anti-integrin β4 (1:250) antibodies. Stained cells were visualized using a Zeiss LSM 510 Meta Confocal Microscope.

### Transwell cell migration and invasion assays

A total of 1.5x10^4^ cells were seeded onto the upper chamber of 24-well Transwell plates. Medium containing 10% FBS was placed in the lower chamber and served as a chemoattractant. Cells were allowed to migrate for 18 hours at 37 °C, the cells on the upper surface of the filter were removed by gently wiping with a cotton swab. The cells that had migrated to the Transwell were fixed and stained with crystal violet. The migrated cells were visualized by a microscope.

For the invasion assay, 1% Matrigel was added to the upper chamber before cell migration, and cells were allowed to migrate for 24 hours at 37 °C.

### Cell adhesion assay

A total of 5x10^3^ cells labeled with cell tracker were plated onto 96-well plates coated with laminin, and the cells were cultured in RPMI-1640 medium with 5% fetal bovine serum at 37 °C in an incubator with 5% CO2 for 3 hours.

### Chromatin immunoprecipitation (ChIP) assay

A total of 1X10^7^ NIH/3T3-MARVELD1-V5 cells were used for the ChIP assay. ChIP was performed as described previously (Weinmann and Farnham, 2002). Promoter regions were PCR amplified with specific pairs of primers that are listed in Table s1.

### Luciferase assay

HTR-8/SVneo cells were cultured in 12-well plates (Corning Glass, Tewksbury, MA, USA) and were transfected with 1 μg of PC or MARVELD1-V5 along with an integrin β4 promotor construct. At 24 hours after transfection, luciferase activity was assayed with 10 μl of lysate using the Dual-Luciferase reporter assay system from Promega (Madison, WI, USA) and a Dynex (Sunnyvale, CA, USA) luminometer. The transfection efficiency was normalized to Renilla luciferase activity. Luciferase activity was measured using the CytoFluorplate 4000 Luminescence Microplate Reader (ABI, Foster City, CA). Three transfection assays were performed to obtain statistically significant data.

### Statistical analysis

Statistical analyses were performed via two-tailed Student’s t-test or one-way ANOVA where indicated. *P*<0.05 was considered as statistically significant, *P*\*<0.05, *P*\*\*<0.01, and *P*\*\*\*<0.001. Data from at least three independent experiments per group were used for evaluation. Values are presented as the mean ± SEM.

## Results

### Generation of MARVELD1 knockout mice via the conditional Cre-loxP targeting system

In humans, the nuclear factor MARVELD1 is a potential tumor suppressor. Our previous study found that MARVELD1 is a novel protein and is downregulated in multiple cancers via methylation of its promoter. More importantly, it mediates cell adhesion and migration by regulating integrin family molecules. To further investigate the role of this novel protein in developmental processes, we generated the MARVELD1 knockout mice using a conditional targeting vector (Fig. 1A). We obtained *MARVELD1*^+/+^, *MARVELD1*^+/−^ and *MARVELD1*^-/-^ mice (Fig. 1B). The *MARVELD1*^-/-^ mice are able to develop into normal adults.

**Figure 1.**
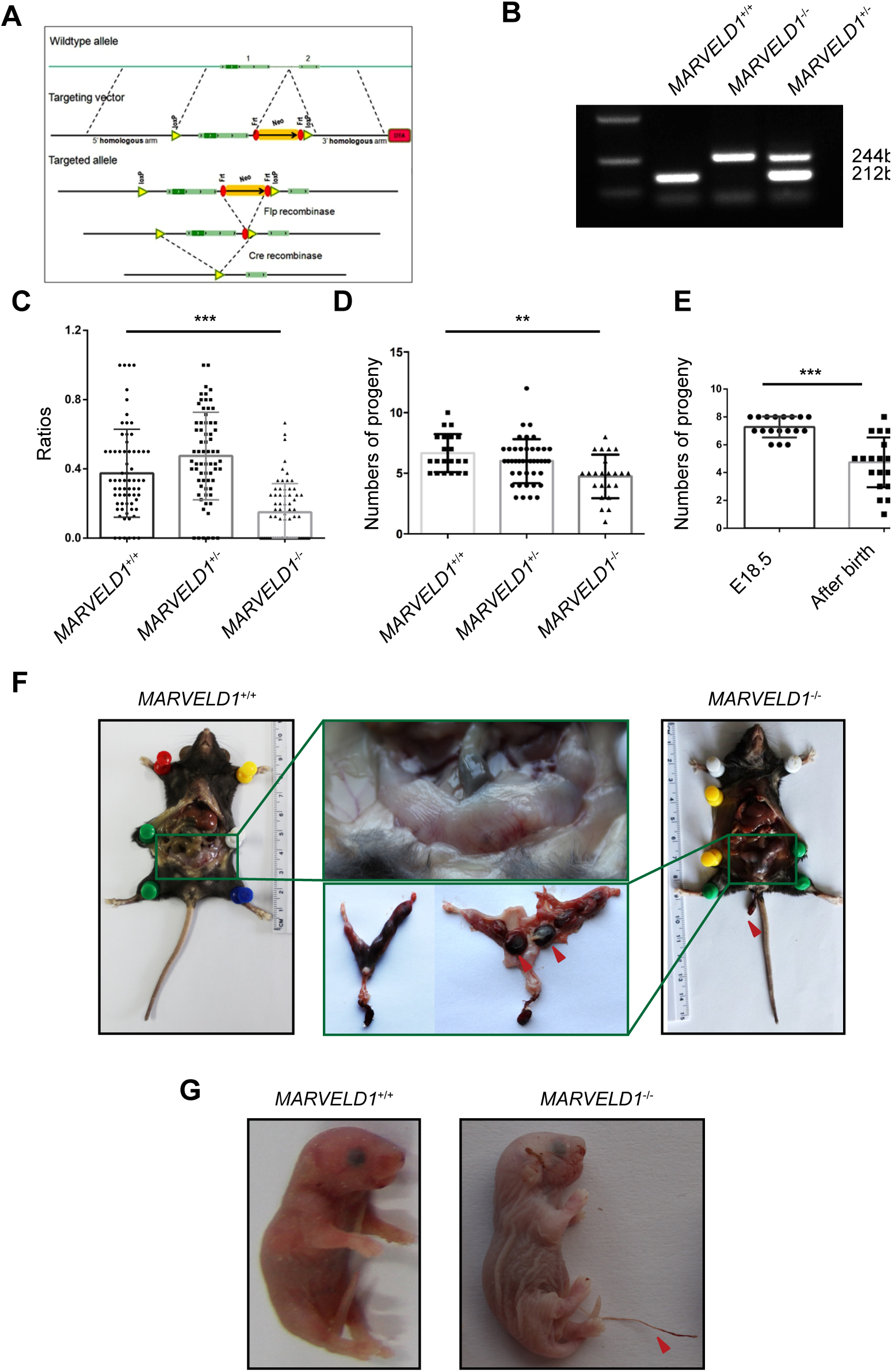
**MARVELD1 knockout mice exhibit birthing difficulties**. (A) Strategy for the modification of the MARVELD1 locus in ES cells and mice. Shown in this figure is a schematic diagram of the targeting vector, the wild-type locus, and the modified MARVELD1 locus before and after excision of the PGK-Neo cassette with Cre recombinase. (B) Genotype analysis of DNA from the tails of offspring animals for the modified MARVELD1 allele. PCR reactions were carried out with primer pairs specific for the wild-type MARVELD1 locus and the MARVELD1 knockout locus from which the neo cassette had been excised by breeding with Cre mice (see text and Materials and Methods for details). The sizes of the expected PCR products are indicated on the right side, and the genotypes are indicated above each lane. (C) Genotype distribution of progeny in *MARVELD1*^+/−^ intercross litters, n=68. (D) Numbers of progeny in *MARVELD1*^+/+^ (n=21), *MARVELD1*^+/−^ (n=42), *MARVELD1*^-/-^ (n=23) intercross litters.(E) Numbers of progeny in *MARVELD1*^-/-^ intercross litters before birth (n=18) and after birth (n=23). (F) Difficult birth phenotype of the *MARVELD1*^-/-^ mother, and the successful production of the wild-type mother. (G) The long umbilical cord phenotype of newborn offspring from the *MARVELD1*^-/-^ intercross.

### MARVELD1 knockout mice have lower newborn offspring rate

The knockout of MARVELD1 does not result in a distinct phenotype after birth. We used *MARVELD1*^-/-^ mice to identify the phenotype of MARVELD1 deletion. At first, we assessed MARVELD1 function by determining the birth rate of *MARVELD1*^+/−^intercrossed mice. The newborn offspring genotypes of 68 litters were counted. Notably, 0.3749 ± 0.03082 were wild-type, 0.4746 ± 0.03068 were heterozygotes, and 0.1505 ± 0.01998 were homozygotes. Compared with the wild-type offspring, the birth rate of homozygotes offspring was significantly lower (*P*<0.001) (Fig. 1C). We also collected *MARVELD1*^+/+^ intercross, *MARVELD1*^+/−^ intercross, and *MARVELD1*^-/-^intercross newborn offspring genotypes to confirm the results. Compared with the wild-type intercross group, the birth rate in the homozygous group was significantly lower (*P*<0.01) (Fig. 1D); this data indicated that the *MARVELD1*^-/-^ mice have lower birth rates than that of wild-type mice. To further identify the reasons that impact the knockout mice birth rate, we analyzed the E18.5 embryo numbers and the newborn offspring pup numbers after intercrossing *MARVELD1*^-/-^ mice; compared with the E18.5 embryo numbers, the newborn offspring pup numbers were significantly lower (P<0.001) (Fig. 1E), which indicated an increased tendency to lose pups before childbirth.

### The pregnant MARVELD1 knockout mice exhibited a placenta accreta phenotype

In addition to decreased pup numbers in the MARVELD1 knockout mice after E18.5, we also found that the placenta attached to the maternal uterus (Fig. 1F) and that the newborn offspring exhibited long umbilical cords (Fig. 1G) by the time of birth. These phenotypes suggested the MARVELD1 knockout mice, to a large extent, exhibited placenta accreta. We further tested the expression levels of MARVELD1 in the E18.5 wild-type mouse organs and found that MARVELD1 was highly expressed in the placenta (Fig. 2A); additionally, the expression levels were increased in the placentas from E10.5 to E18.5 (Fig. 2B), which is the time point that correlates with the trophoblast cell invasion process. We further identified the localization of MARVELD1 in wild-type E18.5 placentas. As shown in Fig. 2C, MARVELD1 was highly expressed in the trophoblast cells, especially the trophoblast cells that tended to migrate into the maternal decidua. These observations indicate that MARVELD1 plays a role in trophoblast cell invasion.

**Figure 2.**
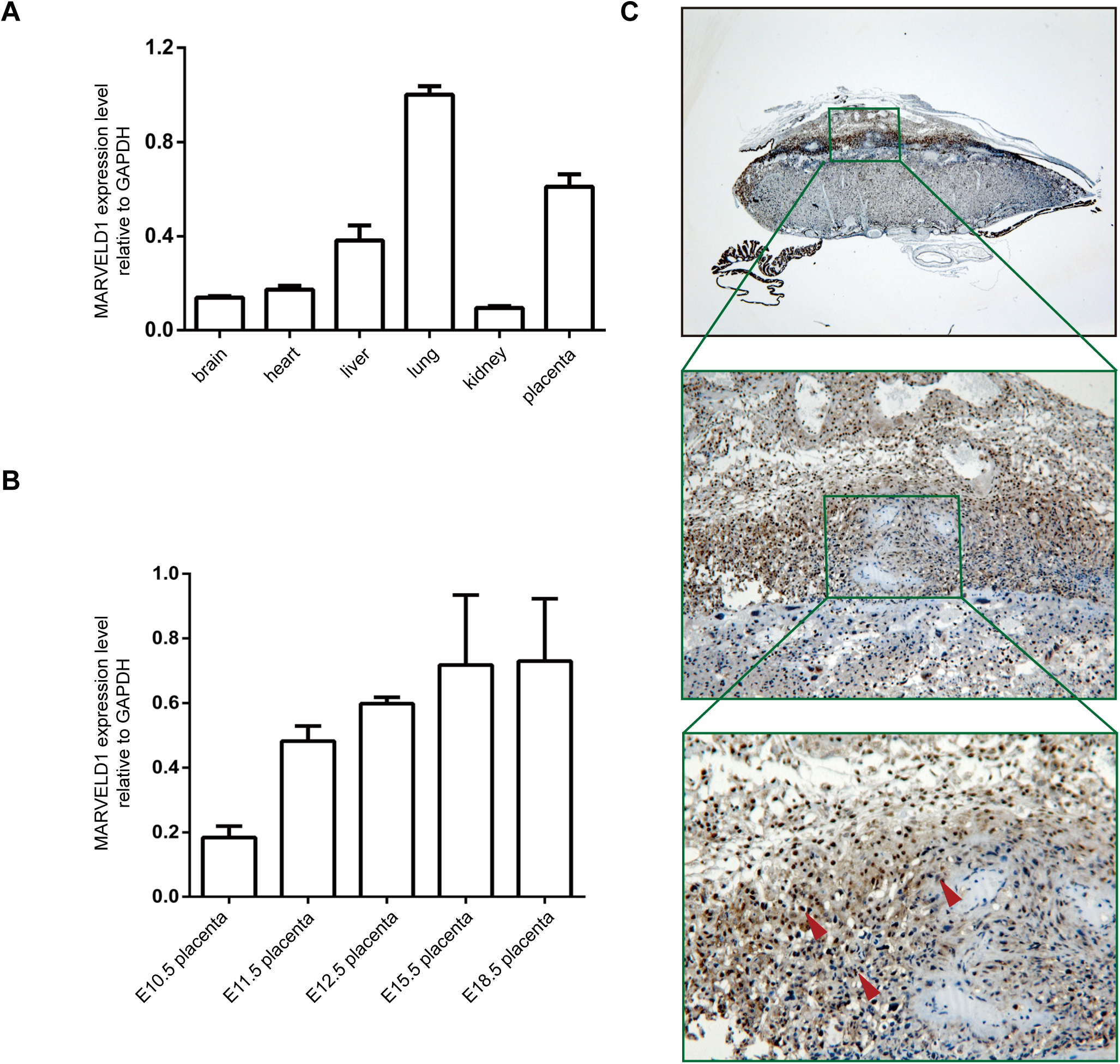
**MARVELD1 is highly expressed in the placenta and invasive trophoblast cells**. (A) Relative expression of the mRNA level encoding MARVELD1 in E18.5 organs, including brain, heart, liver, lung, kidney and placenta, was determined by real-time RT-PCR. GAPDH mRNA was used to normalize the variability in template loading. The data are reported as the mean ± SEM. (B) Relative expression of mRNA encoding MARVELD1 in E10.5, E11.5, E12.5, E15.5 and E18.5 placentas. (C) Immunohistochemical analysis of E18.5 placenta show that MARVELD1 is highly expressed in the placenta and invasive trophoblast cells.

### The trophoblast cells of the MARVELD1 knockout mouse placenta exhibited increased cell invasion

The high expression of MARVELD1 in the placenta and the trophoblast cells partially revealed that MARVELD1 plays a role in trophoblast cell invasion. To further confirm this hypothesis, we investigated trophoblast cell invasion in placenta accreta mice. HE staining showed abnormal adhesion of the placenta to the myometrium (Fig. 3A). Using immunohistochemical staining of Cytokeratin18, a trophoblast cell marker, we observed that the maternal decidua was highly occupied by trophoblast cells in placenta accreta mice (Fig. 3B). After we confirmed the placenta accreta phenotype, the rate of placenta accreta was determined in *MARVELD1*^-/-^ female mice. In total, 45.95% of female *MARVELD1*^-/-^ mice exhibited the placenta accreta phenotype, and 70.59% of those mice were pregnant for the first time (Fig. 3C). To double confirm the over-invasion of trophoblast cells in MARVELD1 knockout placenta, the E18.5 placentas of wild-type mice and MARVELD1 knockout mice were compared. We used HE staining to examine the architecture of the junction layer, which contain the trophoblast cells that invade the maternal decidua. The boundary of the junction layer in wild-type mice was clear; in contrast, the boundary in MARVELD1 knockout mice was indistinct (Fig. 3D). Using immunohistochemical staining, we visualized a clear difference of trophoblast cell invasion between E18.5 wild-type placenta and MARVELD1 knockout placenta. An increase in trophoblast cell invasion from the E18.5 knockout placentas into the maternal decidua was observed (Fig. 3E). These observations confirmed a strong connection between MARVELD1 and the over-invasion of trophoblast cells.

**Figure 3.**
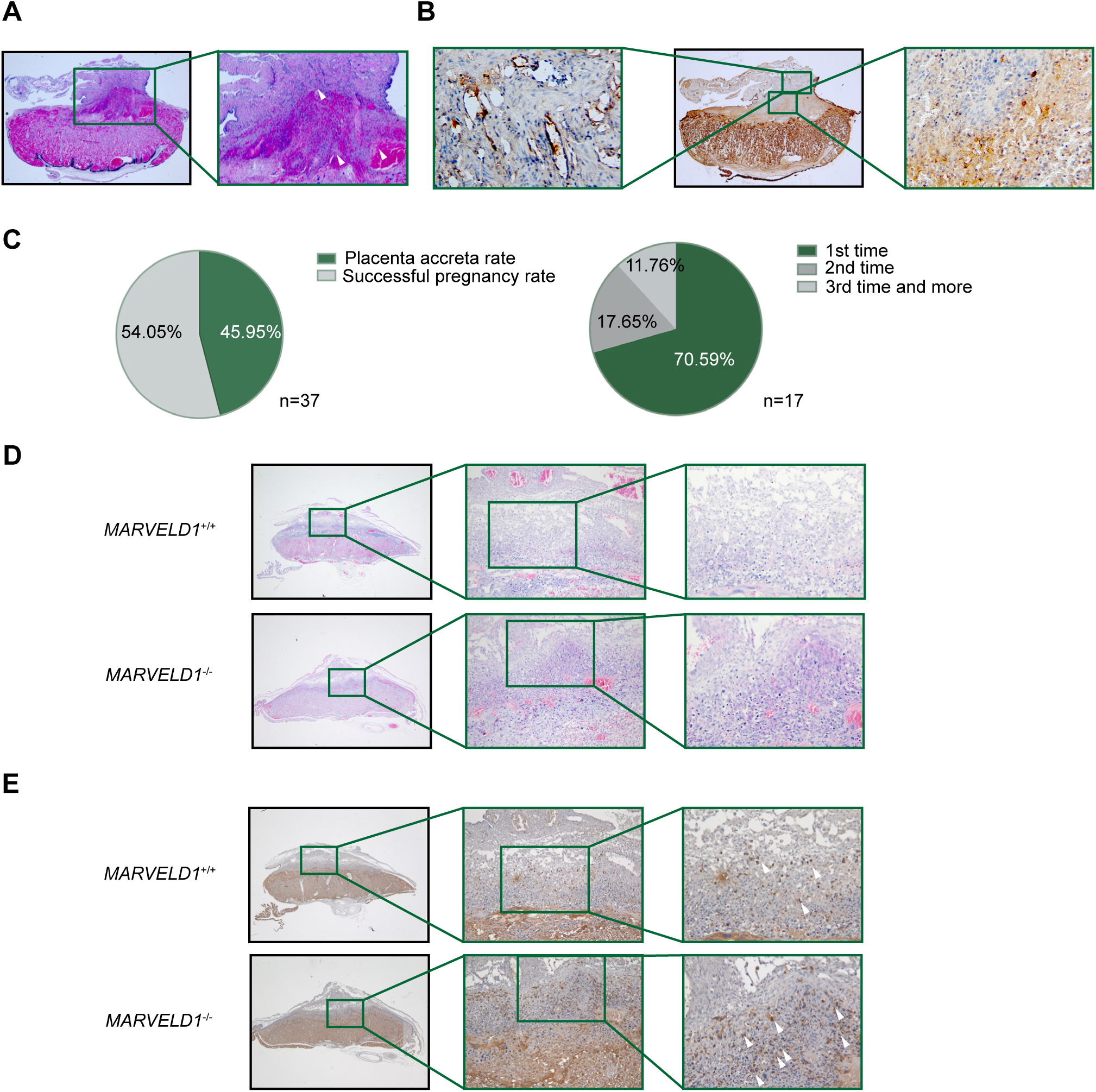
***MARVELD1*^-/-^ placenta exhibits abnormal architecture and trophoblast cell over-invasion** (A) HE staining showing the attachment of the placenta to the maternal uterus from a MARVELD1 KO dam.(B) Immunohistochemistry of placentas from MARVELD1 KO dams for Cytokeratin18, a trophoblast cell marker. (C) The occurrence the dystocia phenotype in *MARVELD1*^-/-^ female mice, n=37; the relationships between the dystocia phenotype and the times of pregnancy in (D) HE staining of the placenta from E18.5 MARVELD1 KO and control dams showing the abnormal architecture of the junctional region in *MARVELD1*^-/-^ placenta. (E) Immunohistochemistry of the placenta from E18.5 *MARVELD1*^-/-^ and control dams for Cytokeratin18. Note the over-invasion of trophoblast cells in the *MARVELD1*^-/-^placenta.

### MARVELD1 was downregulated during trophoblast cell migration and invasion

To further elucidate the contribution of MARVELD1 during trophoblast cell invasion, the HTR8/SVneo cell line was developed as an in vitro model. In human cell lines, the MARVELD1 expression level is higher in mesenchymal cell lines and is lower in epithelial cell lines. The MARVELD1 expression level in the HTR8/SVneo cells was in the mid-range (Fig. 4A). Ten ng/µL TGF-β1 was added to HTR8/SVneo cells, and afterwards, the invasion ability of the HTR-8/SVneo cells was improved (Fig. 4B). Then, we tested the expression of epithelial markers, mesenchymal markers, integrin β1, integrin β4, and MARVELD1. As shown in Fig. 4C, treatment with 10 ng/µL TGF-β1 led to a significant decrease in E-cadherin expression level (*P*<0.001) and an increase in the expression of mesenchymal markers N-cadherin and snail (*P*<0.001). When the invasion ability of HTR8/SVneo cells was improved, we detected a notable increase of integrinβ1 and a significant decrease of integrin β4; MARVELD1 was slightly but not that significantly decreased. These results indicated that the HTR8/SVneo cell line was an ideal model to assess MARVELD1-mediated trophoblast cell invasion.

**Figure 4.**
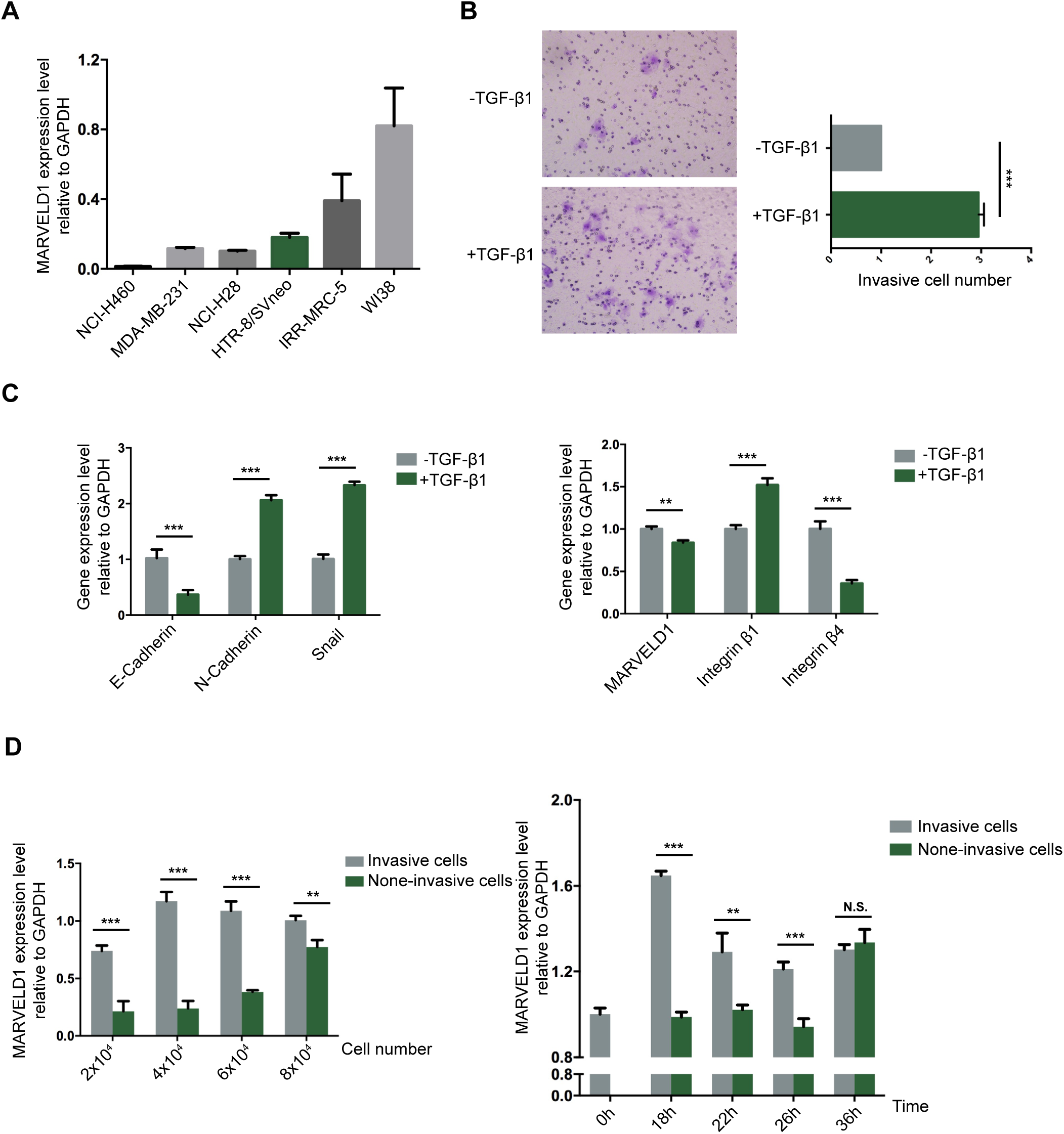
**MARVELD1 and integrin β4 are downregulated in HTR8/SVneo cells during EMT** (A) Relative expression of MARVELD1 mRNA in human cell lines was determined by real-time RT-PCR. GAPDH mRNA was used to normalize the variability in template loading. The data are reported as the mean ± SEM. (B) For the Transwell assay, 10 ng/µL TGF-β1 was added to the cells, and the images were acquired after 24 hours. (C) The mRNA levels of epithelial markers, mesenchymal markers, integrin β1, integrin β4, and MARVELD1 with/without TGF-β1 determined by real-time RT-PCR. (D) The mRNA level of MARVELD1 during cell migration stratified by cell number and migration time. Non-migrated cells above the chamber and migrated cells below the chamber were collected separately.

We further used the Transwell chamber to select cells by their migration ability. Cells above the membrane and below the membrane were collected separately. Different cell numbers were used: 2X10^4^, 4X10^4^, 6X10^4^ and 8X10^4^. MARVELD1 expression in the cells above the membrane and below the membrane was significantly different at each cell density except 8 X10^4^ (Fig. 4D). Different time points were also tested by plating 6X10^4^ cells onto the chamber and collecting cells above the membrane and below the membrane after 0, 18, 22, 26, and 36 hours. The difference in MARVELD1 expression between the migrated cells and the non-migrated cells exhibited the highest difference at 18 hours, and the difference in expression diminished afterwards; there was no significant difference after 36 hours. These data indicate that MARVELD1 is expressed at low levels in the migrated trophoblast cells.

### MARVELD1 regulated trophoblast cell invasion via integrin β4

In human non-small cell lung cancer, we found that MARVELD1 mediated the balance of integrin β1/β4 and the EMT phenotype(Yao et al., 2015). Since integrin β1/β4 belongs to integrin family, which participates in trophoblast cell invasion, we transfected the HTR8/SVneo cells with siRNAs that can downregulate the mRNA expression level of MARVELD1 to assess if MARVELD1 regulates the expression of integrin β1/β4 in trophoblast cells. We speculated that MARVELD1 had the same roles in trophoblast cells as in human non-small cell lung cancer. We found, however, that the roles were not the same, but they were similar. Compared with the scrambled siRNA-transfected HTR8/SVneo cells, MARVELD1 siRNA significantly decreased the mRNA expression level of MARVELD1 (*P*<0.001) as well as the expression level of integrin β4 (*P*<0.001), but the expression level of integrin β1 was not significantly changed (Fig. 5A). The protein level of integrin β1/β4 correlated with the mRNA results (Fig. 5B, 5C). These results indicated that MARVELD1 specifically regulated the expression of integrin β4 in trophoblast cells. To determine whether this relationship between MARVELD1 and integrin β4 was involved in the migration and invasive ability of HTR8/SVneo cells, the cells were transfected with scrambled siRNA, siRNA for MARVELD1, vector, siRNA for MARVELD1 and vector, integrin β4, and siRNAs for MARVELD1 and integrin β4. The migration and invasive abilities were positively correlated, and the abilities from highest to lowest were as follows: siMARVELD1~siMARVELD1 +PC; scrambled siRNA~ scrambled siRNA +PC; integrin β4 +siMARVELD1; and integrin β4. Compared with the scrambled siRNA group, the migration and invasive abilities were enhanced in the cells transfected with siRNA for MARVELD1 (Fig. 5D). These results support the synergy between MARVELD1 and integrin β4 in the invasion of HTR8/SVneo cells.

**Figure 5.**
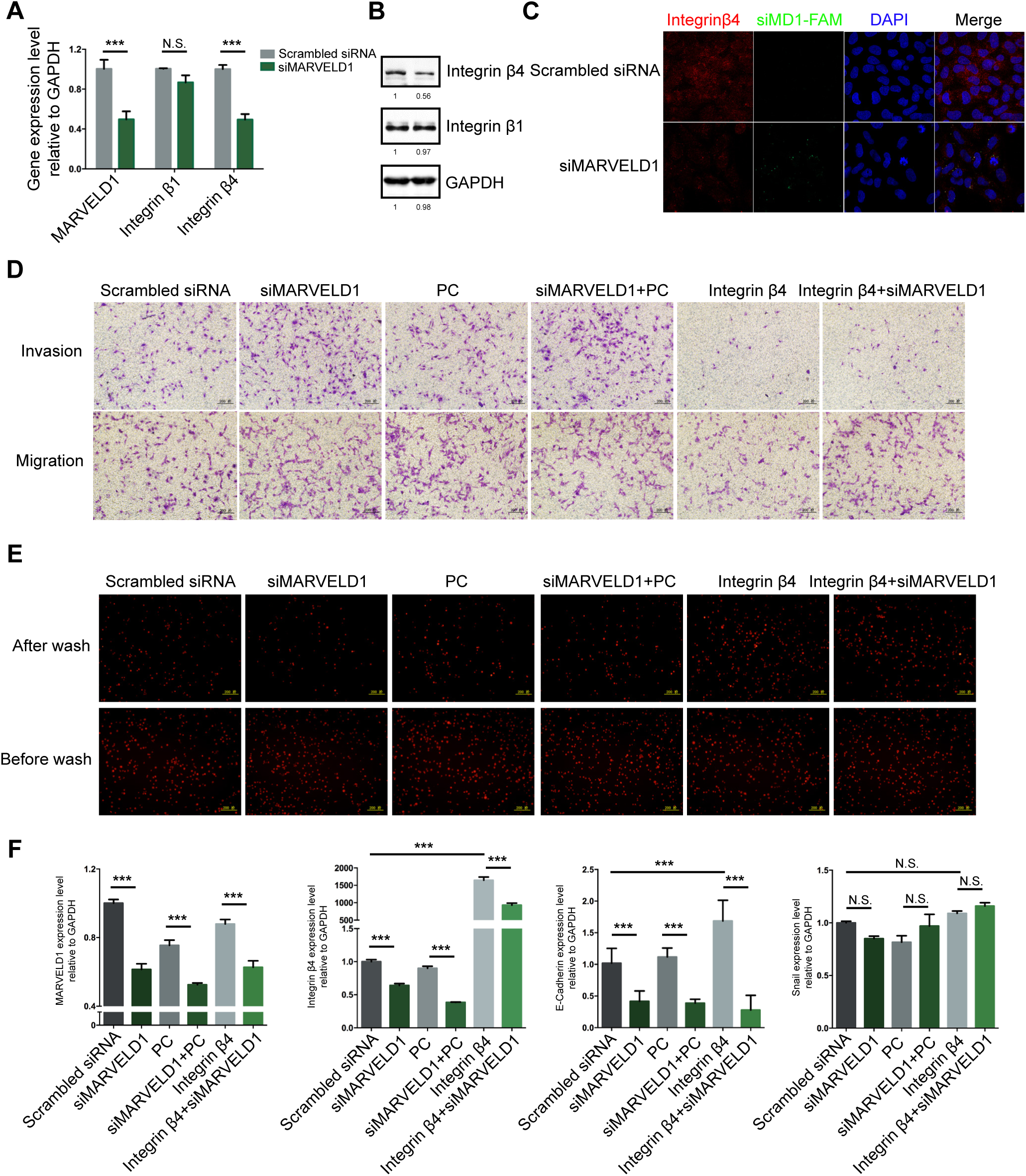
**MARVELD1 regulates trophoblast cell invasion via the integrin β4-mediated cell adhesion pathway** siRNA for MARVELD1 was added to cells to downregulate MARVELD1 mRNA expression levels. (A) Relative expression of integrin β1, integrin β4, and MARVELD1 mRNA in cells was determined by real-time RT-PCR. GAPDH mRNA was used to normalize the variability in template loading. (B) Western blot analysis of the expression of integrin β1 and integrin β4. GAPDH was used as a loading control. (C) Images of cells immunostained with antibodies against integrin β4. (D) Cells were labeled with cell tracker, and cell adhesion ability was identified after 3 hours. (E) Transwell assay to analyze cell migration and invasion. (F) Relative expression of the mRNA level encoding MARVELD1, integrin β4, E-cadherin and N-cadherin in cells. As determined by real-time RT-PCR, GAPDH mRNA was used to normalize the variability in template loading.

Since integrin β4 promoted its signaling events and mediated cell adhesion to ECM, we also tested the adhesion ability of cells transfected with scrambled siRNA, siRNA for MARVELD1, vector, siRNA for MARVELD1 and vector, integrin β4, and siRNAs for MARVELD1 and integrin β4. The adhesion ability of each group was negatively correlated with their migration and invasive abilities (Fig. 5E). These results further support that MARVELD1 regulates trophoblast cell invasion via integrin β4-dependent cell adhesion.

As the invasion process of trophoblast cell is somewhat like an EMT process, EMT markers were also tested to double confirm our hypothesis. As shown is Fig. 5F, the E marker was down regulated when the invasive ability was increased, and there were no significant changes in the M marker. These data fully confirm our previous hypothesis.

### MARVELD1 increases the expression of integrin β4 by enhancing its promoter activity

As the expression of integrin β4 is correlated with MARVELD1, this raises the question of how MARVELD1 regulates the expression of integrin β4. We have demonstrated before that MARVELD1 could specifically bind to the promoter of integrin β4 and activate it, which then enhanced the expression of integrin β4 in human cancer cell lines. To determine whether the same mechanism occurs in mouse cells, NIH/3T3-MARVELD1-V5 cells were used in a ChIP assay. As shown in Fig. 6A, MARVELD1 specifically bound to the integrinβ4 promoter. Luciferase assays were performed in HTR8/SVneo cells transfected with MARVELD1-V5 plasmids. As shown in Fig. 6B, the cells transfected with MARVELD1-V5 resulted in significantly higher integrin β4 promoter activity than PC control cells. These data demonstrate that MARVELD1 regulates the expression of integrin β4 by enhancing its promoter activity.

**Figure 6.**
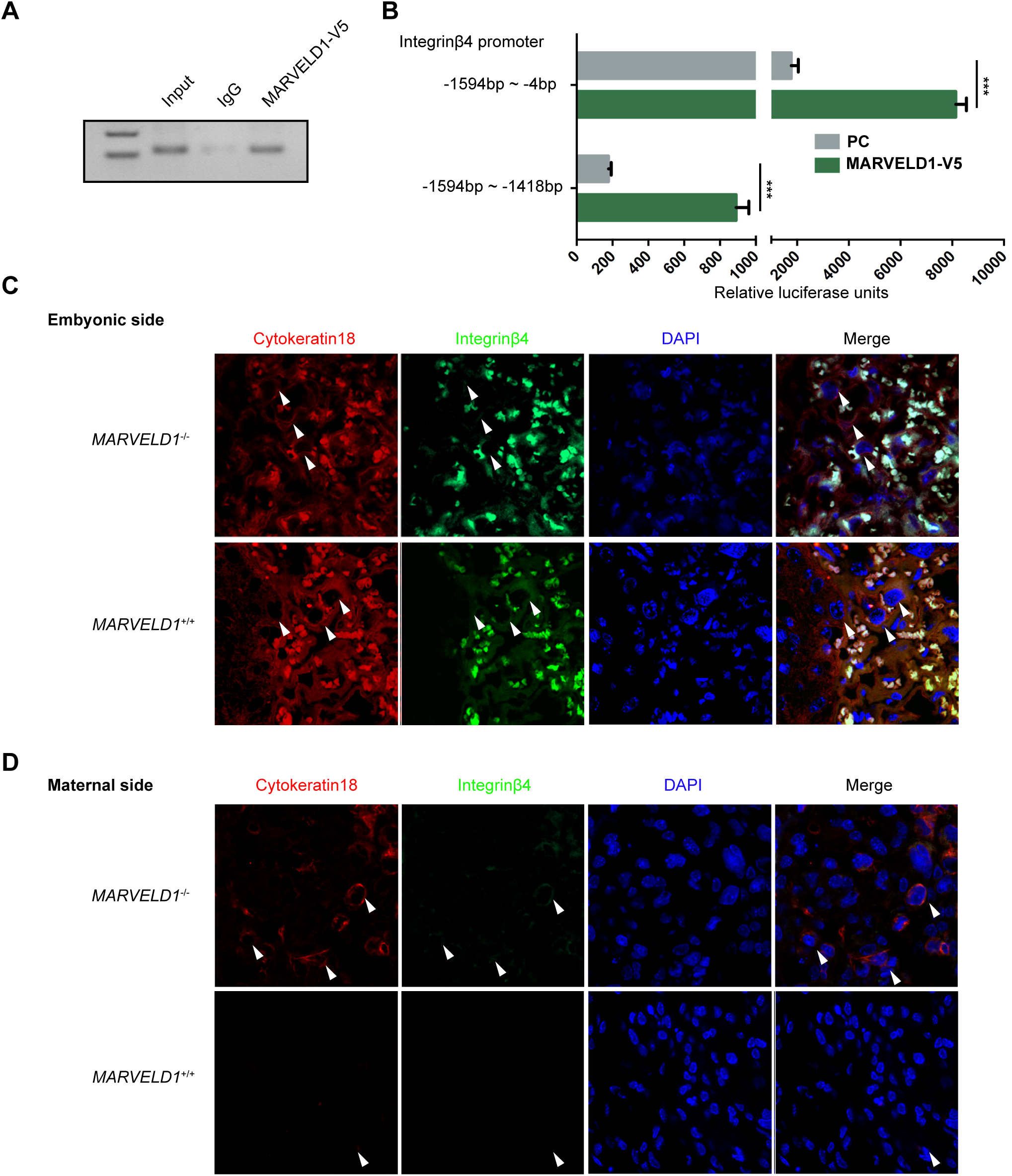
**MARVELD1 increases the expression of integrin β4 by enhancing its promoter activity** (A) A ChIP assay was performed by using HTR8/SVneo cell lysate. Chromatin was immunoprecipitated using a V5-tag specific antibody. PCR was carried out using specific primers for the integrin β4 promoter. (B) Cells were transfected with the integrin β4 (-1,594~-449) or the integrin β4 (-1,594~-1,418) luciferase reporter plasmid. The firefly luciferase activity was measured 24 hours post-transfection and normalized to Renilla luciferase activity. Values represent the mean ± SEM of three independent experiments. (C) Embryonic side: images of E18.5 *MARVELD1*^+/+^ and *MARVELD1*^-/-^ placenta sections immunostained with antibodies against integrin β4 (green) and Cytokeratin18 (red) as well as DAPI staining (blue). (D) Maternal side: images of E18.5 *MARVELD1*^+/+^ and *MARVELD1*^-/-^ placenta sections immunostained with antibodies against integrin β4 (green) and Cytokeratin18 (red) as well as DAPI staining (blue).

### Loss of MARVELD1 affects trophoblast cell invasion via integrin β4 expression changes in vivo

The migration capability of HTR8/SVneo cells could be affected by the decreased expression of MARVELD1 via its effects on the integrin β4-mediated cell adhesion pathway, thus explaining the over-invasiveness of trophoblast cells observed in MARVELD1 knockout mice in vivo. The invasion of trophoblast cells is a precise process that is regulated by complex molecules; therefore, we aimed to further confirm the molecular mechanisms in vivo. In E18.5 placentas, the deletion of MARVELD1 resulted in a decrease of the integrin β4 (Fig. 6C) expression, demonstrating an overall increase in its cell adhesion pathway. In Fig. 6D, we show in E18.5 MARVELD1^-/-^ placentas that there was a large amount of trophoblast cells that invaded into the maternal decidua compared to wild-type mice. The observed effect of MARVELD1 knockout in the E18.5 placenta supports the notion that the deletion of MARVELD1 downregulates the expression of integrin β4 and promotes the over-invasion of trophoblast cells (Fig. 7A).

**Figure 7.**
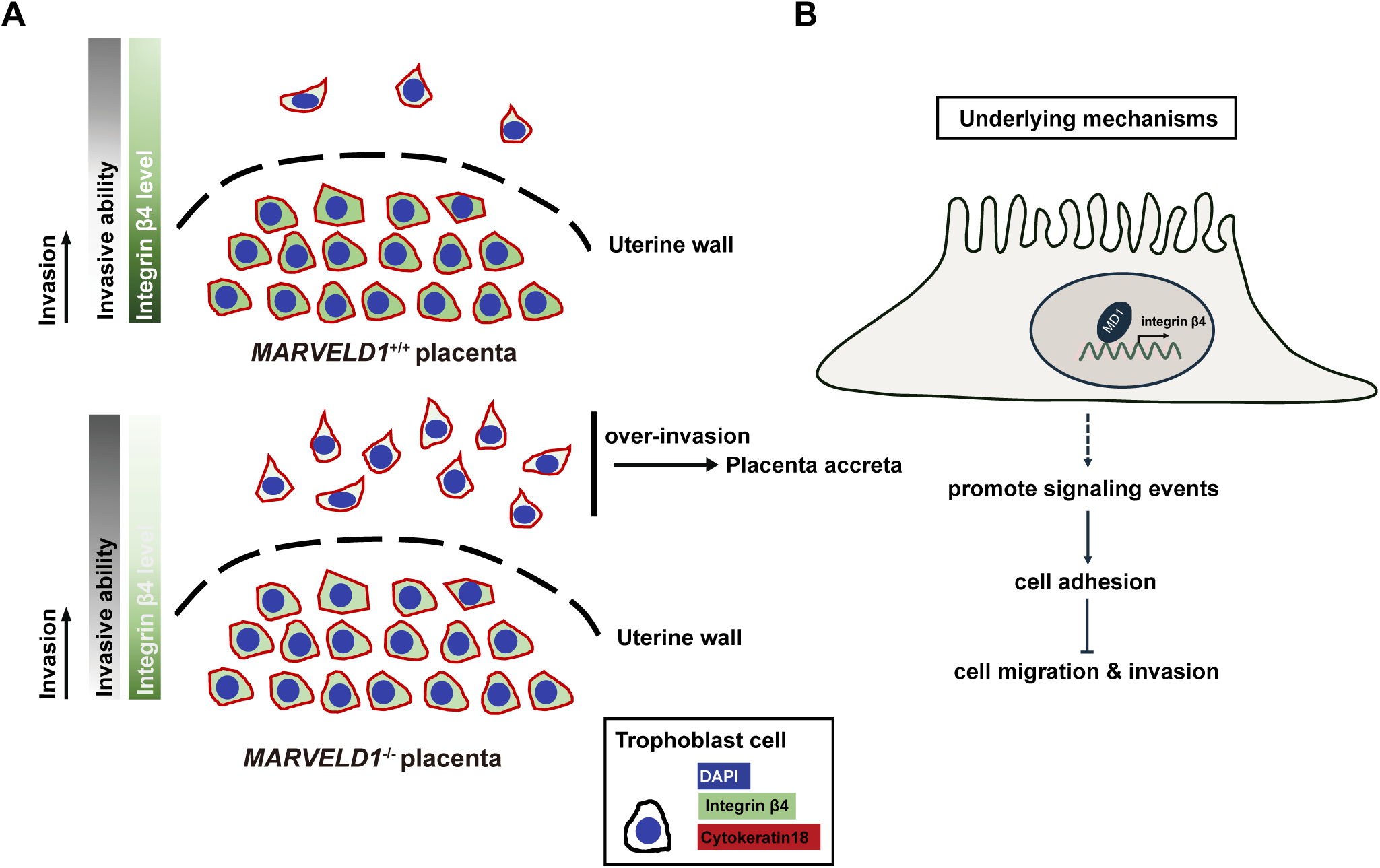
**Proposed models for MARVELD1-mediated trophoblast cell invasion in vitro and in vivo** (A) Working model of MARVELD1 in mediating in vivo trophoblast cell invasion. When MARVELD1 is knocked out, the expression level of integrin β4 is downregulated and the adhesive ability of cells is suppressed, which boost cell migration and invasion, leading to trophoblast cell over-invasion and the placenta accreta phenotype in *MARVELD1*^-/-^ mice. (B) Underlying molecular mechanisms. MARVELD1 binds to the integrin β4 promoter to activate its transcription. Then, integrin β4 upregulates the signaling processes that facilitate cell adhesion and suppress cell migration and invasion.

## Discussion

The function of MARVELD1, a novel nuclear factor that mediates cell adhesion and migration in lung cancer, in mouse development is still unclear. Since gene knockout mice are ideal models in which to study gene function, we developed the MARVELD1 knockout mouse to assess phenotypes after gene deletion, and more importantly, to explore the molecular mechanisms of this novel protein. Our data indicated that the knockout of MARVELD1 caused low birth rates but did not affect the numbers of embryonic offspring. We identified a placenta accreta phenotype, characterized by the attachment of the placenta to the maternal uterus, and the over-invasion of trophoblast cells in MARVELD1 knockout mice. Consistent with these observations, we found that the loss of MARVELD1 suppressed integrin β4 expression and the integrin β4-dependent cell adhesion process, demonstrating that MARVELD1 mediates integrin β4 expression and plays important roles in placenta development, especially in trophoblast cell invasion (Fig. 7A, 7B).

MARVELD1 is a novel MARVEL domain-containing protein that is downregulated in multiple human cancers(Wang et al., 2009). The MARVEL-domain containing proteins such as MYADM(Aranda et al., 2011) and occludin(Huber et al., 2000) are involved in cell adhesion and migration. Our previous studies have revealed that MARVELD1 regulates integrin β1 though a pre-mRNA processing pathway(Wang et al., 2013) and that MARVELD1 is involved in TGF-β1-mediated EMT by regulating the balance of integrin β1 and β4 expression in NSCLC cells(Yao et al., 2015). These findings indicate that MARVELD1 plays a role in cell adhesion and migration in cancers. Since adhesion and migration are basic cell characteristics, MARVELD1 is highly conserved between different species, and therefore, we hypothesized that MARVELD1 plays a role during mouse development.

To test the hypothesis, MARVELD1 knockout mice were developed. The *MARVELD1*^-/-^ mice can develop into normal adults but have lower birth rates compared to wild-type mice. During the birthing process in MARVELD1 knockout mice, we visualized that the placentas were difficult to detach from the maternal uterus due to the over-invasion of trophoblast cells. Cell migration is central to multiple physiological processes, including embryonic development and cancer metastasis. During both development and normal homeostasis, cells interact with each other and the surrounding environment, including morphogen and the ECM, to maintain tissue structure, organization and function(Paluch et al., 2016). During embryonic development, cells migrate through interacting with their surrounding environment; the tissue remodeling and morphogenesis processes involve the migration of different cell types (Muthuswamy and Xue, 2012). The migration of cells engaged in tissue-remodeling events contribute to cell invasion during development. Rodents and human have similar hemochorial placenta, and trophoblast cell invasion plays important roles in this type of placenta. The trophoblast cell invades into the maternal uterus in search of spiral arterioles and veins. Trophoblast cells breach the spiral arterioles and replace the endothelial and smooth muscle cells (Red-Horse et al., 2006). These processes are important for both proper fetal perfusion and attachment(Maltepe and Fisher, 2015). Previous studies have indicated that the trophoblast cell invasion process is similar to epithelial-to-endothelial transition(Maltepe and Fisher, 2015); E-cadherin and integrin α6β4 are downregulated during trophoblast cell invasion(Maltepe and Fisher, 2015).

Here, we identified that the downregulation of MARVELD1 decreases the expression of integrin β4 and integrin β4-dependent cell adhesion process both in vivo and vitro, which correlates with the phenotype observed in MARVELD1 knockout mice. These in vivo and in vitro studies further elucidated the contribution of MARVELD1 in placenta development. Notably, previously reported findings mainly highlighted integrin β4 as an epithelial marker, and its expression was an indicator for EMT. Based on our results, we suggest that integrin β4 is also an EMT initiator, and this process is also under the control of MARVELD1 during placenta development. Our study suggests a sequence of events that result in the over-invasion of trophoblast cells; additional studies are required to further illustrate this process. Although a lot has been discovered about the placenta in recent decades, there is a lot more regarding placental abnormalities and pregnancy complications that remain to be discovered. In summary, our study has elucidated a novel function of the MARVELD1 gene in trophoblast cell invasion.

## Acknowledgements

The HTR8/SVneo cell line was a gift from Prof. Yongjun Yang (Jilin University). This work was supported by the National Natural Science Foundation of China No. 31571323 and Shenzhen Municipal Basic Science Foundation No. JCYJ20140417173156097 to YL; the China Postdoctoral Science Foundation No. 2015M571426 to YC.

### Author contributions

YL and YC conceived the idea, designed experiments, provided the conceptual framework for the study, and wrote the manuscript. HZ further contributed to manuscript preparation. YC, HZ, FH, LY, and CQ performed the experiments. YC, HZ, YZ and WL analyzed the data, PD contributed to conceptual evaluation of the project.

